# Combined Antibiotic and Herbicide Pollution Accelerates the Horizontal Transfer of Antibiotic Resistance Genes in Coastal Microbial Communities

**DOI:** 10.64898/2026.02.27.708170

**Authors:** Hui Yang, Hongyue Ma, Yang Yang, Shaoguo Ru, Yifan Liu, Meiru Wang, Hui Miao, Ziyi Guo, Liqiang Yang, Pengfei Cui

## Abstract

Graphical abstract

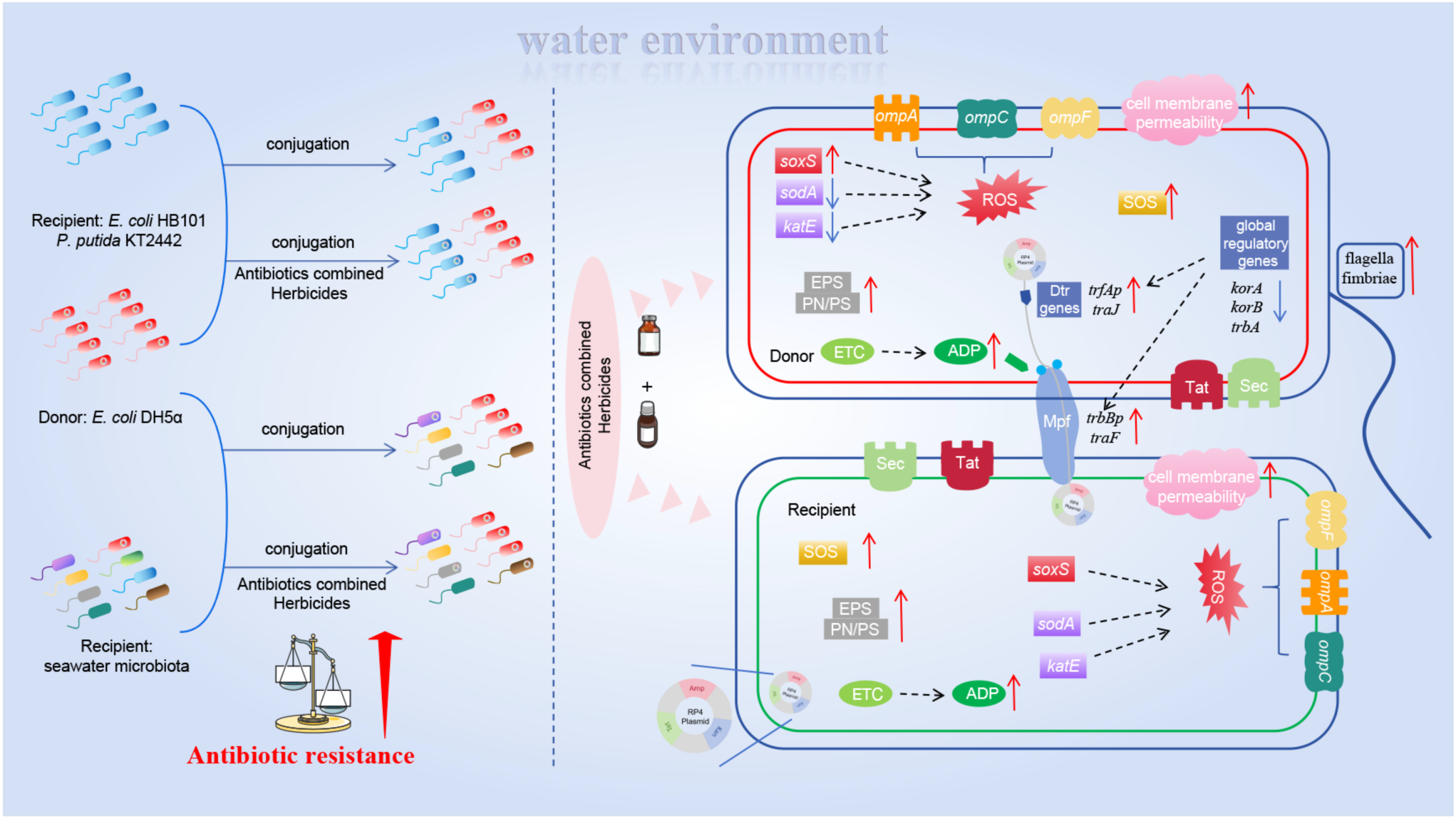

The extensive use of herbicides and antibiotics in aquaculture has led to compounded pollution of coastal waters, marked by the co-occurrence of herbicides, antibiotic residues, and antibiotic resistance genes (ARGs). Plasmid-mediated conjugative transfer is a major driver of the dissemination and evolution of ARGs, yet the influence of herbicides alone or in combination with antibiotics on this process in aquatic bacterial communities remains unclear. In this study, we demonstrated that triazine herbicides, alone and in combination with a fluoroquinolone antibiotic, significantly promote the conjugative transfer of the broad-host-range RP4 plasmid. Using *Escherichia coli* DH5α as a donor, we observed increased plasmid transfer to multiple recipients, including *E. coli* HB101, *Pseudomonas putida* KT2440, and a natural coastal seawater microbial community. The enhanced transfer under co-exposure was associated with several interacting mechanisms, including elevated intracellular reactive oxygen species (ROS) and adenosine triphosphate (ATP) levels, increased cell membrane permeability, altered extracellular polymeric substance (EPS) composition, and upregulated expression of conjugation-related genes. Overall, this work underscores the potential role of combined herbicide and antibiotic contamination in shaping the microbial resistome of coastal ecosystems and provides insights to inform strategies for mitigating the spread of antibiotic resistance.

## 1. Introduction

Antibiotic resistance driven by antibiotic resistance genes (ARGs) presents a serious global health threat, potentially causing up to 10 million deaths annually by 2050 (O’Neill, 2014). The presence of ARGs extends far beyond clinical settings and was recognized by the United Nations Environment Programme (2017) as one of six emerging environmental issues of global concern. A large-scale survey detected 2,561 ARGs across six distinct habitats, conferring resistance to 24 classes of antibiotics. These ARGs were not only present in both human and environmental but also highly mobile, with 23.78% posing direct health risks to humans (Zhang et al., 2022).

Horizontal gene transfer (HGT) is the primary mechanism for ARGs dissemination, occurring via transduction, natural transformation, and conjugation (Tan et al., 2019; Errington et al., 2001). Among these, plasmid-mediated conjugative transfer is recognized as the most efficient route for ARGs propagation among environmental microorganisms (von Wintersdorff et al., 2016). Notably, RP4 multidrug resistance plasmids can transfer to wastewater microbiota and have a broad host range, covering over 13 bacterial phyla and 46% of genera (Huang et al., 2022).

Traditionally, antibiotics have been considered the main drivers of plasmid-mediated conjugation. For example, enrofloxacin, a widely used fluoroquinolone, promotes conjugative transfer of the fluoroquinolone resistance gene *qnrS* (Zhao et al., 2022a). However, emerging evidence suggests that non-antibiotic pollutants, such as ketoprofen (Zhang et al., 2024a), fungicide (Zhang et al., 2023), quaternary ammonium biocides (Hu et al., 2024) and phenolic compounds (Ma et al., 2021), can also modulate ARGs transfer. In aquatic ecosystems—especially coastal zones—multiple pollutants often co-occur, such as enrofloxacin from aquaculture effluents and triazine herbicides from agricultural runoff (Yang et al., 2019). Yet, little is known about the combined effects of triazine herbicides and enrofloxacin on plasmid-mediated ARGs conjugation.

Triazine herbicides disrupt the D1 protein of photosystem II (PSII) in plants and are frequently introduced into marine environments through surface runoff (Zhang et al., 2024b). They have been shown to impair reproductive and endocrine functions in aquatic organisms (Wang et al., 2022) and to reduce marine primary productivity, disrupt global carbon cycling, and potentially drive marine desertification (Yang et al., 2024). Moreover, triazines can exert antibiotic-like effects by inducing bacterial oxidative stress and altering gut microbiota composition (Liu et al., 2019; Stara et al., 2013; Zhang et al., 2012; Zhao et al., 2022b).

Known mechanisms by which environmental pollutants promote conjugative transfer include increased ROS production, elevated cell membrane permeability, and upregulation of SOS and conjugation gene (Zhang et al., 2023). While triazines are known to induce oxidative stress, it remains unclear whether this ROS-mediated pathway contributes significantly to ARGs conjugation. Furthermore, microorganisms under environmental stress often regulate extracellular polymeric substances (EPS) to improve survival. For example, enrofloxacin can stimulate EPS secretion in algae, increasing cell surface hydrophobicity and adhesion (Cheng et al., 2023). Furthermore, adenosine triphosphate (ATP) is essential for the construction of membrane-translocation systems required during plasmid transfer (Chen et al., 2005), may also be affected under combined pollutant exposure, yet this relationship remains unexplored.

Therefore, the present study aimed to elucidate the effects of triazine herbicides—individually and in combination with enrofloxacin on the conjugative transfer of the RP4 plasmid. Three experimental models were established to assess plasmid transfer from *Escherichia coli* DH5α to intrageneric (*E. coli* HB101), intergeneric (*Pseudomonas putida* KT2442), and natural seawater bacterial communities. Potential mechanisms were explored through phenotypic and genotypic analyses, including intracellular ROS and ATP levels, membrane permeability, EPS composition, conjugation gene expression, and transcriptomic profiling. These findings provide new insights into the complex impacts of co-pollution on ARGs dissemination in marine environments and offer a theoretical basis for developing integrated control strategies to mitigate ARGs spread.

## 2. Materials and methods

### 2.1. Bacterial Strains, Plasmid, Antibiotics, and Triazine Herbicides

Three distinct conjugation models were established to investigate the transfer dynamics of ARGs. The donor strain for all experiments was *Escherichia coli* DH5α carrying the conjugative RP4 plasmid, which encodes resistance to kanamycin (Km), tetracycline (Tet), and ampicillin (Amp). **Model 1**: *E. coli* HB101 (streptomycin-resistant) served as the intrageneric recipient strain. **Model 2**: *Pseudomonas putida* KT2440 (chloramphenicol-resistant) served as the intergeneric recipient strain. **Model 3**: Natural seawater microbiota collected from Tuan Island Bay, Qingdao, China, served as the environmental recipient community (see Text S1 and S2).

Four triazine herbicides were tested: atrazine (Yuanye Biotechnology, China), terbutryn (MedChemExpress, China), prometryn and ametryn (both from Aladdin Biochemical Technology, China). Enrofloxacin was purchased from Sigma-Aldrich (USA), and all other antibiotics from Solarbio (China).

### 2.2. Growth kinetics determination

Growth kinetics of donor *E. coli* DH5α and recipient *E. coli* HB101 were assessed under varying concentrations of enrofloxacin (0, 0.5, 5, 50, 500 μg/L) and atrazine (0, 5, 50, 500, 5000, 50000 μg/L). Bacterial suspensions were adjusted to 10^5^ CFU/mL in LB medium containing the respective treatments. Aliquots (150 μL) were transferred into 96-well plates and incubated at 37 °C for 8 h. To assess the growth progress, the absorbance at a wavelength of 595 nm was measured at hourly intervals using a multifunctional microplate reader (Thermo Fisher Scientific, China). Each condition was tested with at least three biological replicates.

### 2.3. Conjugation experiments

Our study aimed to investigate the influence of triazine herbicides, both individually and in combination with enrofloxacin, on conjugation transfer within aquatic environments. For this purpose, we used liquid media in our conjugation experiments rather than solid media. Our experimental setup followed a methodology similar to a prior investigation (Liu et al., 2021). Donor and recipient strains were washed three times with PBS and adjusted to 10^8^ CFU/mL. Equal volumes were mixed, and enrofloxacin (5 μg/L) was added, followed by triazine herbicides at 5, 50, or 500 μg/L. Cultures were incubated at 28°C for 8 h.

For Models 1 and 2, transconjugants (NT) and recipients (NR) were spread on selective LB agar containing the corresponding antibiotics. Transfer frequency was calculated as NT/NR, and transconjugants was confirmed via plasmid extraction, PCR, and agarose gel electrophoresis (Text S3, Table S2). For Model-3, in accordance with the methodology outlined in a previous study by (Ma et al., 2021), conjugation was evaluated by quantifying changes in the *E. coli*-specific gene *groEL*, the conjugative gene *traF*, and ARGs (*tetA* and *aphA*) from the RP4 plasmid. There were three biological replicates at least per experiment.

### 2.4 EPS extraction and analysis

EPS were extracted from donor and recipient strains using a heat extraction method (Liao et al., 2019). The quantification of polysaccharides was accomplished through the phenol-sulfuric acid method. Furthermore, the protein content was assessed using the BCA Protein Assay Kit (Solarbio, China). There were three biological replicates at least per experiment.

### 2.5 Detection of ROS and cell membrane permeability

2’,7’-Dichlorofluorescein diacetate (DCFH-DA) were used for measurements of ROS formation. In detail, bacterial suspensions (10^6^ CFU/mL) were incubated with 10 μM DCFH-DA for a duration of 30 minutes at 37°C in a dark environment. Subsequently, unbound probes were removed by washing. Enrofloxacin and the tested triazine herbicides were then introduced and treated as previously described, also in the absence of light. The resulting reaction mixtures were measured fluorescence intensity using a multifunctional microplate reader (Thermo Fisher Scientific, China), the wavelengths were 488 nm (excitation) and 525 nm (emission). There were three biological replicates at least per experiment.

Cell membrane permeability was assessed in accordance with a methodology established in prior research (Yu et al., 2021). Bacterial suspensions were stained with 20 μM of propidium iodide (PI) and incubated at 37°C for 15 min in the dark. Subsequently, all samples were analyzed using a flow cytometer (Beckman FC500-MPL, USA) with the wavelengths of 488 nm (excitation) and 561 nm (emission). There were three biological replicates at least per experiment.

### 2.6 Measurement bacteria ATP content

Conjugation mixtures were washed twice with PBS, and intracellular ATP was quantified using the Enhanced ATP Assay Kit (Beyotime, China).

### 2.7 Analysis of mRNA expression of genes

Total RNA was extracted from the conjugation systems using the Bacteria RNA Extraction Kit (Vazyme, China). Subsequently, this RNA was reverse - transcribed into complementary DNA (cDNA) using HiScript Ⅲ RT SuperMix for qPCR (Vazyme, China). Real-time quantitative polymerase chain reaction (qPCR) was performed on a MasterCycler® ep RealPlex4 (Eppendorf) using Taq Pro Universal SYBR qPCR Master Mix (Vazyme, China). Genes analyzed included oxidative stress (*soxS*), conjugation-related (*korA*, *korB*, *trbA*, *trfAp*, *trbBp*, *traJ*, *traF*), outer membrane (*ompA*, *ompC*, *ompF*), and antioxidant (*sodA*, *katE*). The 16S rRNA served as the reference gene (Yang et al., 2022) (Table S1). There were three biological replicates at least per experiment.

### 2.8 RNA sequencing, PacBio microbial diversity sequencing and bioinformatics analysis

For the RNA sequencing analysis, intergeneric conjugation systems were treated for 2 h with control, 5 μg/L enrofloxacin, 500 μg/L atrazine, and both combined. For PacBio sequencing, seawater microbiota conjugation systems were treated for 8 h with control, 5 μg/L enrofloxacin, 50 μg/L atrazine, and both combined. The bacterial solution was washed three times and dewatered, subsequently rapid freezing with liquid nitrogen. These samples were subsequently preserved at a temperature of −80°C. Then all samples were submitted to Majorbio Co., Ltd. (Shanghai, China). The raw data was analysis according to previous research (Liao et al., 2019).

### 2.9 Statistical Analysis

Data were analyzed using GraphPad Prism 9. To assess significant differences among adjacent columns of distinct colors, we applied an independent-sample *t*-test. Additionally, significant differences among individual columns of the same color were evaluated using a one-way analysis of variance (ANOVA). Statistical significance was denoted as follows: \**p* < 0.05, \*\**p* < 0.01, and \*\*\**p* < 0.001. The data were presented as the mean ± standard deviation (SD).

## 3. Results and discussion

### 3.1. Effects of herbicides alone and in combination with enrofloxacin on conjugative transfer in pure cultures

To explore how the combination of triazine herbicides and enrofloxacin affects conjugative transfer, we conducted a series of conjugation experiments. Prior to these assays, we evaluated the impact of various concentrations of enrofloxacin and atrazine on the growth of both the donor (*E. coli* DH5α) and recipient (*E. coli* HB101) strains. Based on these results (Figure S1), we selected 5 μg/L enrofloxacin and 5, 50, and 500 μg/L triazine herbicides for further analysis. These concentrations were chosen to minimize any confounding effects due to changes in bacterial density. These concentrations preclude the influence of alterations in bacterial abundance on conjugation transfer.

In Model-1, the conjugative frequencies of RP4 increased with escalating concentrations of the four tested triazine herbicides. This effect was more pronounced when enrofloxacin was added. For instance, in the presence of 500 μg/L atrazine, prometryn, and terbutryn combined with enrofloxacin, the transfer frequencies increased by 3.33-fold, 8.55-fold, and 4.31-fold, respectively (Figure 1a). Notably, no continuous increase in transfer frequency was observed in samples treated with 500 μg/L ametryn, and the frequency decreased when combined with enrofloxacin (Figure 1a), which may be attributed to the excessive structural damage caused by high concentration of ametryn to the bacterial cells. In Model-2, the combined exposure to 5 μg/L enrofloxacin and 500 μg/L atrazine resulted in a remarkable 14-fold increase in conjugative frequency (Figure 1b), surpassing the effects of other pollutants in previous studies (Liao et al., 2019; Yu et al., 2021). Subsequently, we found by plasmid extraction, PCR (*traG* gene) and agarose gel electrophoresis experiments that all the transconjugants contained the same plasmid type as the donors, while no plasmid was detected in the recipients, which confirmed the successful transfer of the RP4 plasmid (Figure S2.).

**Figure 1.**
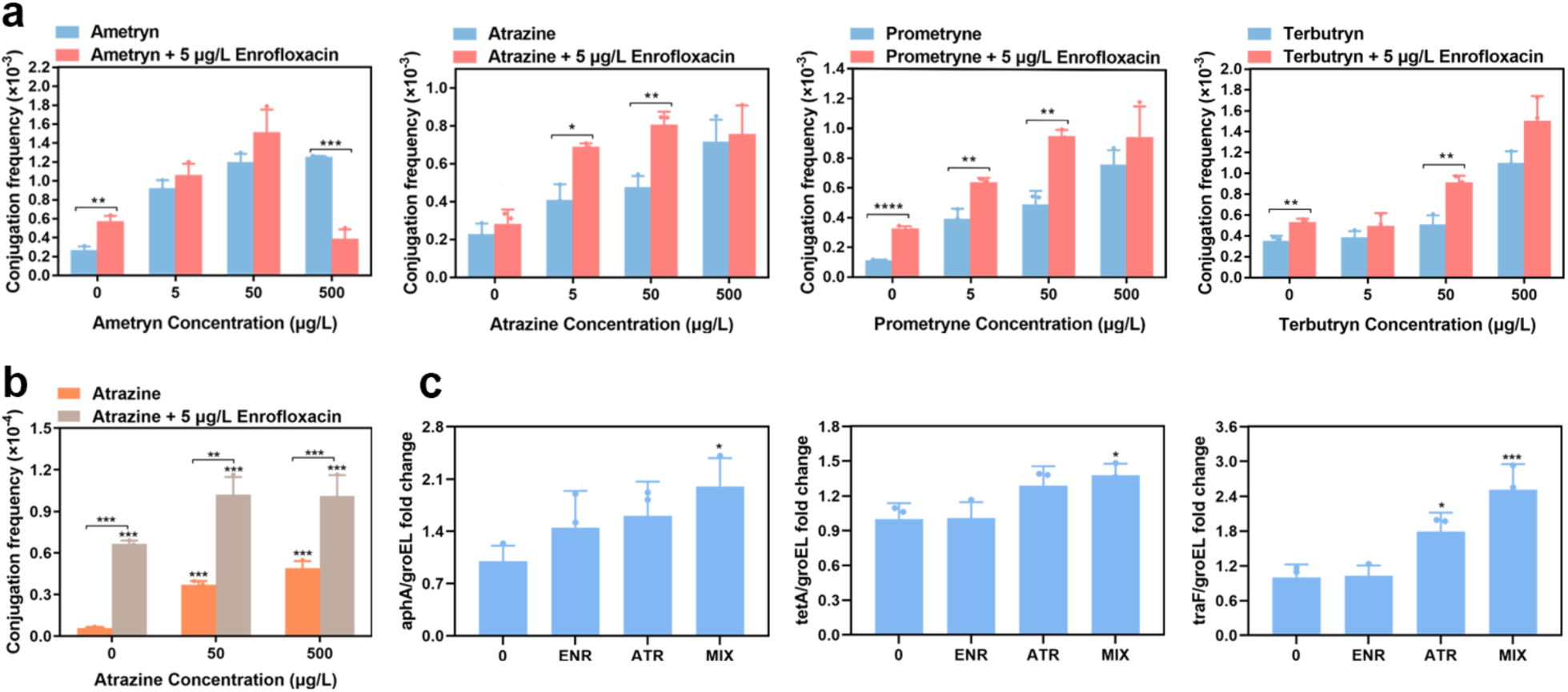
Effect of triazine herbicides (ametryn, atrazine, prometryn, terbutryn) alone and combined with enrofloxacin, on conjugative transfer frequency. (a) Intragenus conjugation system. (b) Intergenus conjugation system. (c) Ratios of RP4 genes to an *E. coli*-specific gene (*groEL*) under atrazine (ATR), enrofloxacin (ENR), and combined (MIX) exposure in the seawater conjugation model.

These findings suggest that both atrazine and enrofloxacin, individually and in combination, can promote conjugative transfer of ARGs within and across bacterial genera. And intra-genus conjugation frequency was generally one order of magnitude higher than inter-genus transfer. This indicates that the co-occurrence of triazine herbicides and enrofloxacin in aquatic environments may contribute significantly to the spread of ARGs across diverse microbial communities. To elucidate the mechanism underlying the promotion of RP4 plasmid conjugative transfer by atrazine alone or in combination with enrofloxacin, we conducted a series of exploratory experiments.

### 3.2. Influences in the conjugative transfer in seawater indigenous microbiome and transconjugants analysis

The co-occurrence of triazine herbicides and enrofloxacin in seawater has been documented, but whether this co-contamination affects the dissemination of ARGs in natural microbial communities remains unclear. To address this, we established a relatively intricate conjugative transfer model using *E. coli* DH5α carrying the RP4 plasmid as the donor and a bacterial community extracted from fresh seawater as the recipient. This design more closely mimics natural environmental conditions compared with experiments using pure cultures.

In this system, the RP4 plasmid was introduced into the seawater bacterial community via donor *E. coli*. Conjugative transfer was assessed by monitoring the ratio of plasmid-bearing bacteria to donor bacteria. Specifically, the RP4 plasmid gene *traF* was used as a marker for plasmid abundance, while the *E. coli* chromosomal gene *groEL* represented donor cell numbers, as described previously (Ma et al., 2021). An increased *traF*/*groEL* ratio indicated plasmid transfer from *E. coli* to other members of the community. Our results revealed that, under co-exposure to 5 μg/L enrofloxacin and 50 μg/L atrazine, the *traF*/*groEL* ratio was significantly higher than in the control, showing a 2.51-fold increase (Figure 1c). We also quantitatively analyzed the ARGs (*tetA* and *aphA*) located on the RP4 plasmid using the same method. The ratios of *tetA*/*groEL* and *aphA*/*groEL* were similar to *traF*/*groEL* trends (Figure 1c). These findings indicated that the RP4 plasmid was able to infiltrate the seawater microbiota under combined herbicide and antibiotic exposure.

In this system, a total of 16 transconjugants were isolated from all samples, including Proteobacteria, Bacteroidetes and Firmicutes. In comparison to the control group, two additional species—*unclassified Pseudoalteromonas* and *Shewanella marisflavi*—were detected under co-exposure to atrazine and enrofloxacin (Table S3). Both belong to Proteobacteria, a group of Gram-negative bacteria whose relatively thin cell walls may facilitate plasmid uptake. Moreover, members of Proteobacteria are known to stabilize RP4 plasmids, which could further enhance the spread of antibiotic resistance genes. These findings therefore suggest that co-exposure can broaden the host range of RP4 plasmids, particularly within Proteobacteria.

Notably, several of the transconjugants identified, such as *Acinetobacter johnsonii*, *Shewanella algae* and *Vibrio alginolyticus*, are pathogenic bacteria associated with human health risks (Bergogne-Bérézin and Towner, 1996; Holt et al., 2005; Wang et al., 2021). However, it should be acknowledged that culture-based methods may underestimate the true diversity of RP4 plasmid hosts in natural seawater. Therefore, these findings indicate that co-exposure can expand the host range of the RP4 plasmid, particularly within the phylum Proteobacteria, and poses a considerable health risk.

### 3.3. Effects of Combined Exposure to Atrazine and Enrofloxacin on Conjugation Gene Expression and SOS Response

Conjugative transfer of the RP4 plasmid involves multiple key regulatory systems, including global regulators, mating pair formation (Mpf) genes, and DNA transfer and replication (Dtr) system genes (Zatyka et al., 2001; Zhang et al., 2018). In our study, we aimed to investigate whether co-exposure to enrofloxacin and atrazine could enhance conjugative transfer by modulating the expression of these regulatory genes, such as the global regulators (*korA*, *korB*, *trbA*), Mpf genes (*trbBp*, *traF*), and Dtr genes (*trfAp*, *traJ*).

Global regulatory genes are known to play a crucial role in inhibiting plasmid transfer and replication (Konig et al., 2009). In our study, we observed that the expression of *korA*, *korB*, and *trbA* was down-regulated to varying degrees in response to both atrazine and combined enrofloxacin treatment (Figure 2a). The Mpf system plays a pivotal role in bringing donor and recipient cells into close proximity during conjugation, facilitating cell membrane fusion and the formation of a junction bridge that serves as an effective transfer channel for DNA (Christie et al., 2014; Zatyka et al., 2001). Our results demonstrated that the co-exposure of enrofloxacin and atrazine promoted conjugative transfer by increasing the expression levels of *trbBp* and *traF* genes (Figure 2b) (Cen et al., 2020; Zhang et al., 2018).

**Figure 2.**
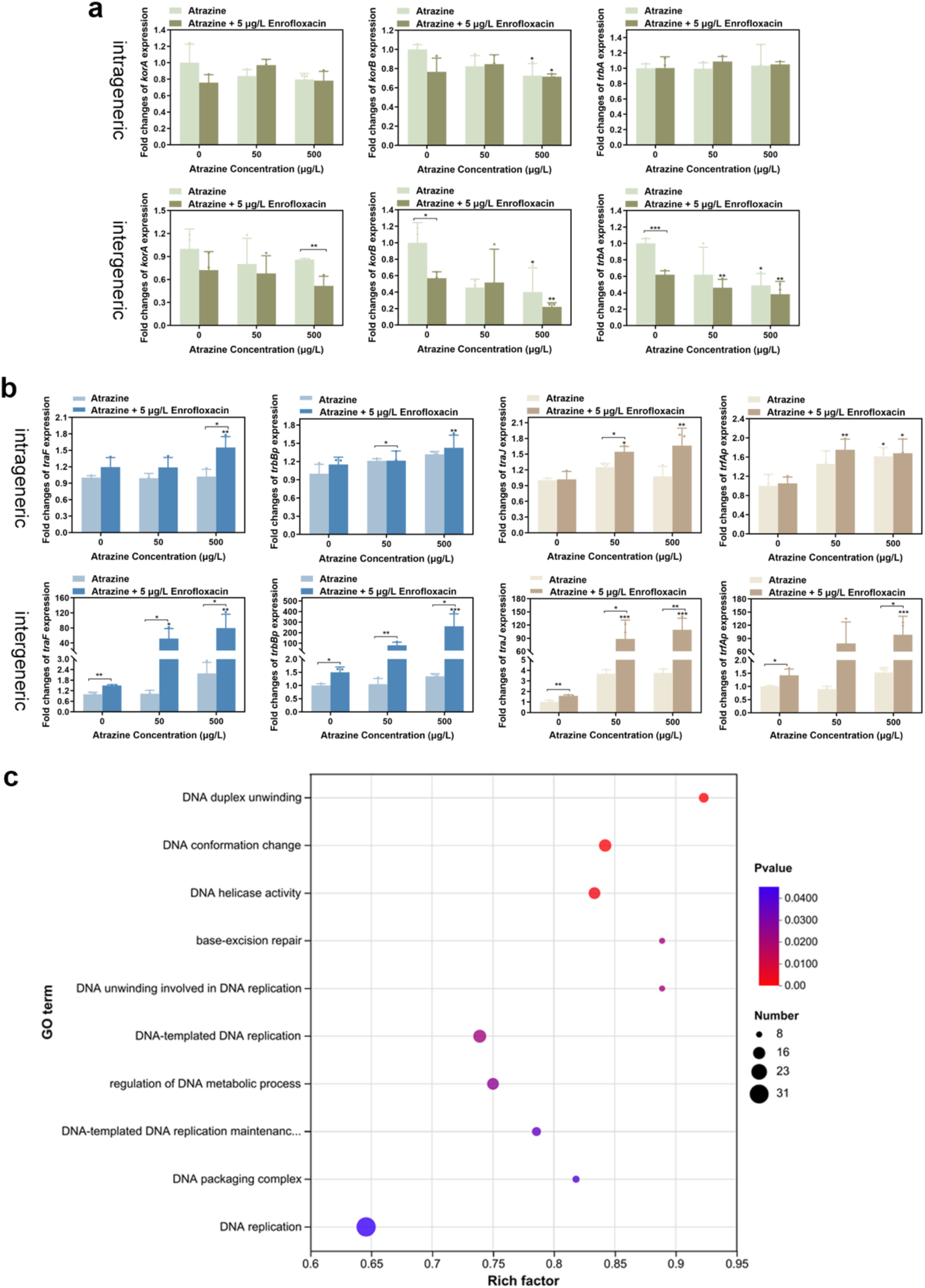
Expression of conjugation genes of *korA*, *korB*, *trbA*. (a) *trfAp*, *traJ*, *trbBp* and *traF*. (b) Under the exposure atrazine alone and in combination with enrofloxacin. (c) GO enrichment analysis of up-regulated SOS response genes in *E. coli* DH5α.

Under the control of the Dtr system, the plasmid undergoes processing into a single strand, which is then transported across the cell membrane from the donor to the recipient bacteria (Samuels et al., 2000; Zatyka et al., 1997). Previous studies have highlighted the importance of *trfAp* as a strong promoter of plasmid RP4 conjugative transfer, and *traJ* as a key role in relaxosome formation (Yang et al., 2022). Upon exposure to atrazine alone or in combination with enrofloxacin, we observed an up-regulation of *trfAp* and *traJ* genes to varying degrees (Figure 2b). These results indicate that combined exposure to enrofloxacin and atrazine can promote the formation of conjugative transfer channels, as well as the mobilization and transfer of plasmids, by inhibiting the expression of global regulatory genes, ultimately increasing the frequency of conjugative transfer.

The SOS response is a cellular stress mechanism activated by abnormal single-stranded DNA or DNA damage, leading to the induction of DNA repair pathways such as homologous recombination and nucleotide excision repair. (Baharoglu et al., 2010; Gao et al., 2023). During conjugative transfer, plasmids can be transmitted in the form of single-stranded DNA, potentially triggering the SOS response (Chen et al., 2005). Moreover, the conjugative transfer process necessitates the induction of the SOS response to facilitate the formation of the second plasmid strand in both donor and recipient cells. Therefore, we conducted an analysis of the transcriptome data, as shown in Figure 2c. SOS response (such as base-excision repair) genes and DNA replication, DNA duplex unwinding and other DNA processing related genes in *E. coli* DH5α were enriched into GO terms, which showed that the enhanced conjugative transfer might be relevant with these processes (Gutiérrez et al., 2021; Quiñones et al., 1989).

### 3.4. The combination of atrazine and enrofloxacin accelerates the generation of energy to facilitate conjugative transfer

ATP plays an essential role in DNA transport during conjugation, as it powers the transmembrane transfer of DNA (Chen et al., 2005). To better understand how energy generation might influence conjugative transfer, we measured ATP levels in both intra- and inter-generic conjugation systems under control and co-exposure treatments. Our data revealed that under the combined exposure of 5 μg/L enrofloxacin and 500 μg/L atrazine, the ATP content in both intra-genera and cross-genera systems exhibited an upward trend, with a significant enhancement of 1.50-fold and 1.77-fold, respectively (Figure 3a).

**Figure 3.**
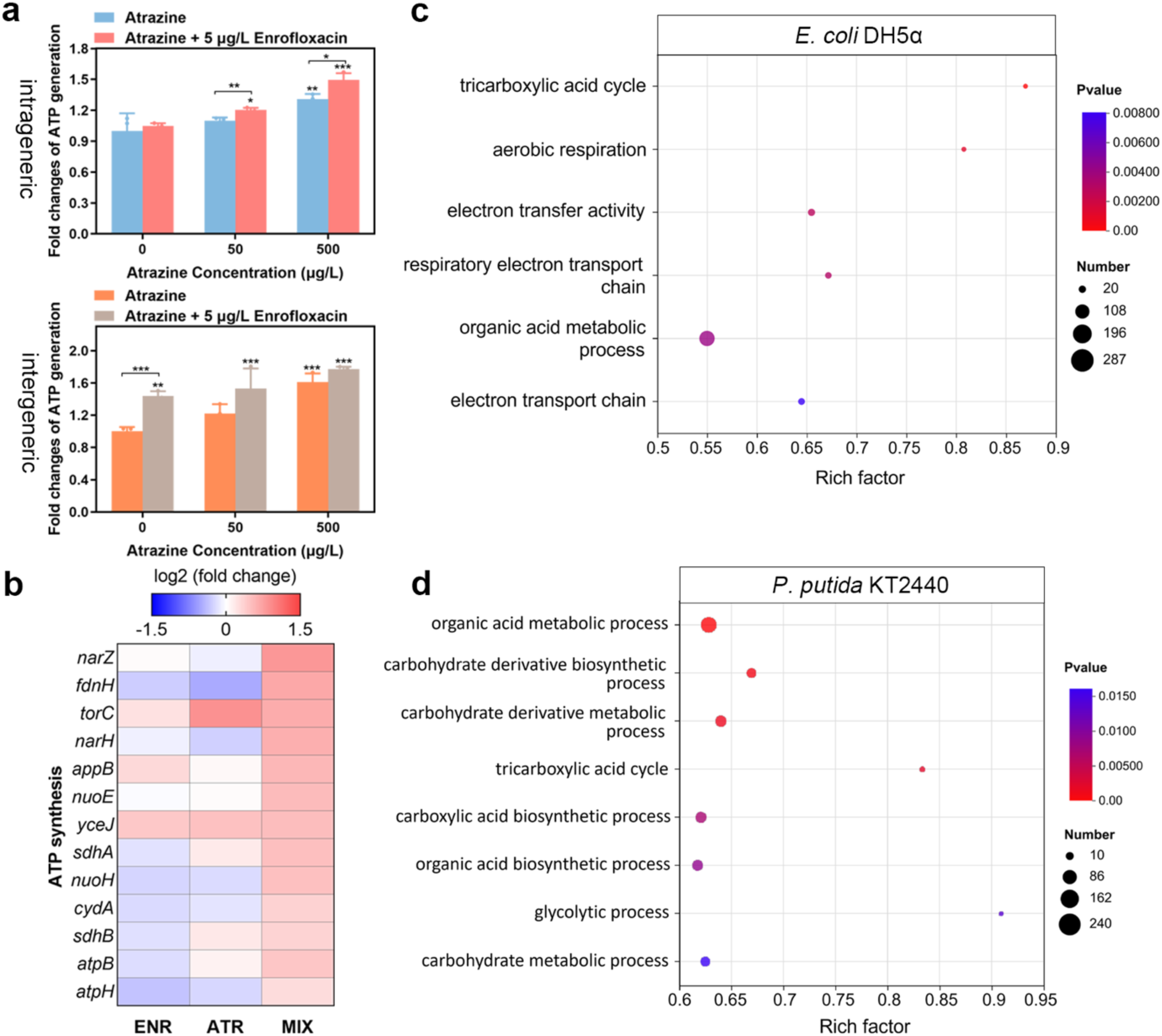
(a) ATP content within conjugation system under exposure atrazine alone or in combination with enrofloxacin. (b) Effects of the expression of ATP synthesis coding-genes under atrazine (ATR), enrofloxacin (ENR), and their combination (MIX) in the donor strain *E. coli* DH5α. (c-d) GO enrichment analysis of up-regulated ATP synthesis genes.

ATP is primarily generated through the electron transport chain (ETC), which comprises transmembrane protein complexes (I-IV), electron transporters like ubiquinone and cytochrome c. The ADP produced by ETC is phosphorylated into ATP through complex V (ATP synthase) (Yu et al., 2020; Zhao et al., 2019). In light of this, we conducted transcriptome analysis. Our data indicated that the transcription levels of ETC and ATP synthase genes increased in response to co-exposure in *E. coli* DH5α (Figure 3b). Meanwhile, glycolytic and tricarboxylic acid cycle and other ATP synthesis genes in *E. coli* DH5α and *P. putida* KT2440 were enriched into GO terms (Figure 3c-d). These findings suggest that augmenting the production of ATP available energy can enhance the frequency of conjugation transfer (Ji et al., 2022; Lu et al., 2018).

### 3.5. Changes of EPS content and cells adhesion in the combination of atrazine and enrofloxacin

Extracellular polymeric substances (EPS) play a critical role in bacterial adhesion and intercellular communication, both of which are essential for conjugative transfer (Liao et al., 2019; Yu et al., 2020). EPS primarily comprise proteins (PN), polysaccharides (PS), DNA, and lipids (More et al., 2014), with polysaccharides and proteins being the major constituents, accounting for 75-89% of the EPS composition (Tsuneda et al., 2003). Therefore, we hypothesize that the concurrent exposure to enrofloxacin and atrazine may affect conjugation transfer by altering the composition of EPS in both donor and recipient cells.

Our findings indicate that the PN/PS ratios of *E. coli* DH5α and *E. coli* HB101 exhibited significant increases under atrazine exposure, especially when combined with enrofloxacin. Conversely, *P. putida* KT2440 exhibited a slight reduction in the PN/PS ratio at low concentrations, but a marked increase at higher concentrations of combined treatment (Figure 4a). Previous research has suggested that enhanced protein synthesis and secretion could lead to an increased PN/PS ratio and improved cell adhesion (Liao et al., 2019). In addition, because polysaccharides contain many hydrophilic groups, a reduction in polysaccharide content within EPS may increase bacterial surface hydrophobicity, which in turn promotes intercellular contact (Li et al., 2022). In line with these mechanisms, our results demonstrated that atrazine, either alone or in combination with enrofloxacin, altered the PN/PS ratio of donor and recipient strains. These findings support the view that co-exposure can modulate cell adhesion by regulating the relative production of polysaccharides and proteins. Furthermore, gene expression analyses confirmed that EPS modifications contribute to changes in bacterial adhesion under co-exposure conditions.

**Figure 4.**
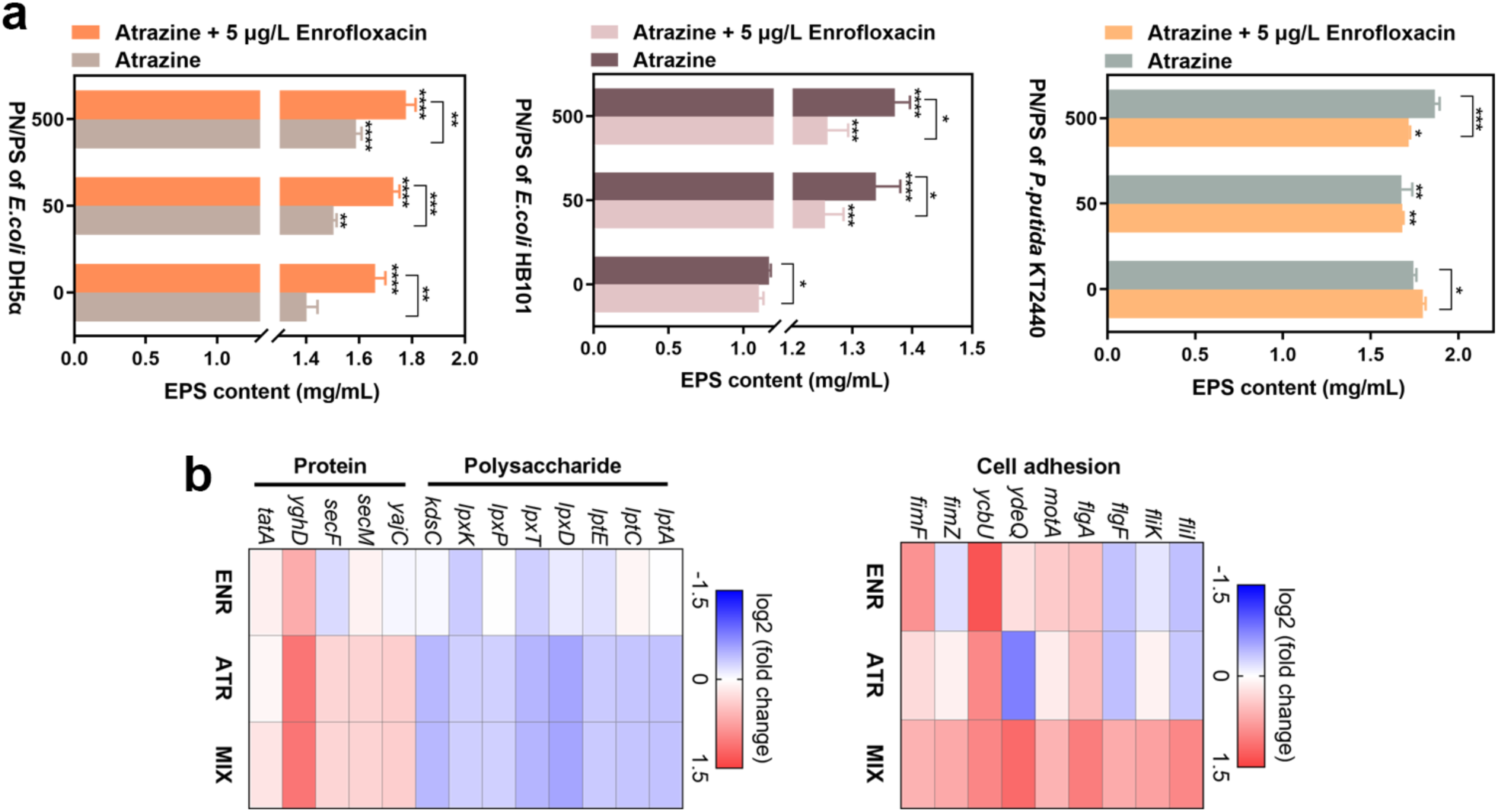
(a) Effects of atrazine alone or combined with enrofloxacin on PN/PS in *E. coli* DH5α, *E. coli* HB101, and *P. putida* KT2440. (b) The changes of the polysaccharide and protein synthesis and export-coding genes/flagella and fimbriae-coding genes expression in *E. coli* DH5α.

Protein secretion pathways were also affected. The Twin-arginine translocation (TAT) and general secretory (Sec) systems are responsible for exporting folded and unfolded proteins, respectively (Crane and Randall, 2017; Palmer and Berks, 2012; Tsirigotaki et al., 2017). The transcriptional analysis revealed notable changes in gene expression for *E. coli* DH5α. Specifically, genes associated with the Sec system (*yajC*, *secM*, *secF*) exhibited a significant upregulation, ranging from 1.26 to 1.30 folds, while the TAT system coding gene (*tatA*) showed an upregulation of 1.15 folds under combined exposure to 5 μg/L enrofloxacin and 500 μg/L atrazine (Figure 4b). Similarly, in *P. putida* KT2440, Sec-related genes (*gspG*, *secD*) and the TAT gene *tatB* were upregulated by 1.26–1.47-fold and 1.29-fold, respectively. In contrast, a downregulation was observed in the expression of genes associated with polysaccharide synthesis and export under the combined exposure to 5 μg/L enrofloxacin and 500 μg/L atrazine. For example, in *E. coli* DH5α (Figure 4b), the expression of genes such as *lptA*, *lptC*, *lpxD*, *lpxT*, and *kdsC* decreased by 0.69-0.83-fold. Similarly, in *P. putida* KT2440, genes including *M8003_05790*, *M8003_16555*, and *lpxC* exhibited reduced expression by 0.65-0.86-fold. Furthermore, structural components that facilitate adhesion were also affected. Genes related to flagella (*fliI*, *fliK*, *flgF*, *flgA*, and *motA*) and fimbriae (*ydeQ*, *ycbU*, *fimZ*, *fimF*) were significantly upregulated in *E. coli* DH5α under combined exposure. Flagella and fimbriae are known to enhance surface attachment and intercellular contact (Avalos Vizcarra et al., 2016; Horstmann et al., 2020). Collectively, these findings suggest that co-exposure to enrofloxacin and atrazine can enhance intercellular contact between bacteria through the modulation of the PN/PS ratio and the promotion of adhesive flagella and fimbriae, thereby playing a pivotal role in the conjugative transfer of the RP4 plasmid.

### 3.6. Increasing of cell membrane permeability in the combination of atrazine and enrofloxacin

Previous research has emphasized the crucial role of the cell membrane as a barrier to the horizontal transfer of ARGs (Garvey et al., 1985; Lu et al., 2018; Thomas and Nielsen, 2005). Within a certain range, increased cell membrane permeability can weaken this obstructive function, thereby promoting the transfer of ARGs between bacteria (Pu et al., 2021; Qiu et al., 2012; Wang et al., 2020a). In our study, flow cytometry analysis showed that the combined treatment of atrazine and enrofloxacin led to a higher percentage of cells stained with propidium iodide (PI), indicating increased membrane permeability in both donor and recipient cells compared to the control group (Figure 5a). Consistent with this, gene expression analyses provided additional evidence for membrane perturbation. Bacterial outer membrane proteins (OMPs) play an important role in the uptake and export of substances from the environments, contributing to membrane permeability and plasmid transfer. The up-regulation of core OMPs genes (*ompA*, *ompC*, *ompF*) in the conjugation system under exposure to 500 μg/L atrazine and 5 μg/L enrofloxacin supports this conclusion (Figure 4b). Transcriptomic analysis further showed that genes associated with the transmembrane transporter complex, cell shape regulation, and other membrane-related transport functions in *E. coli* DH5α and *P. putida* KT2440 were significantly enriched in GO terms (Figure S3).

**Figure 5.**
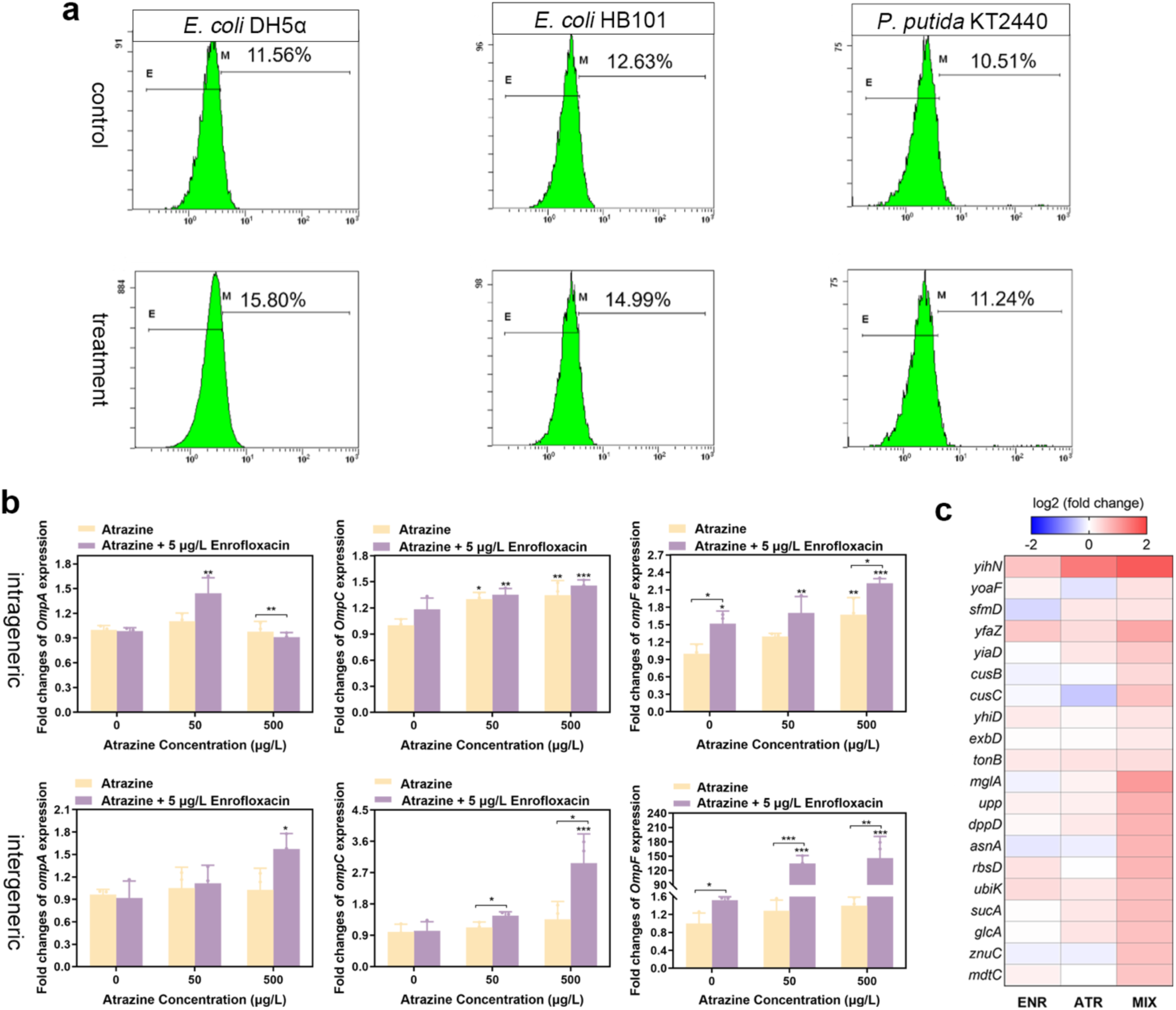
(a) Proportion of PI positive staining cells in control group and combined group. (b) Fold changes in the expression of membrane permeability related gene (*ompC, ompA, ompF*). (c) Effects of the expression of coding-genes related to cell membrane under atrazine (ATR), enrofloxacin (ENR), and their combination (MIX) in *E. coli* DH5α.

Specifically, genes encoding outer membrane proteins (*yoaF*, *yfaZ*, *yiaD*, *sfmD*) and an inner membrane protein (*yhiD*) were upregulated in *E. coli* DH5α (Liao et al., 2021, 2019; Wang et al., 2019). Moreover, genes coding for membrane fusion proteins (*cusB* and *cusC*) (Su et al., 2009; Zhang et al., 2019) and outer membrane transport proteins (*exbD* and *tonB*) (Ollis et al., 2009; Silale and Van Den Berg, 2023) were also significantly upregulated (Figure 5c). Taken together, these results suggest that the combined exposure to atrazine and enrofloxacin increases cell membrane permeability through the upregulation of multiple membrane-related genes, thereby likely facilitating conjugative transfer.

### 3.7 Formation of ROS induced by the combination of atrazine and enrofloxacin

In response to environmental stress, excessive ROS can be generated at the cell membrane, which in turn increases membrane permeability (Mittler, 2002; Qiu et al., 2012). Triazine herbicides, similar to other environmental pollutants, have been reported to induce oxidative stress in bacteria (Zhang et al., 2012). Therefore, in this study we examined intracellular ROS formation and the expression of antioxidant system genes under combined exposure to atrazine and enrofloxacin.

The results showed that ROS production was significantly elevated under co-exposure. Specifically, treatment with 500 μg/L atrazine and 5 μg/L enrofloxacin resulted in a 1.38-to 1.50-fold increase compared with the control (Figure 6a). Consistently, expression of oxidative stress-related genes was upregulated. For example, in *E. coli* DH5α, the *iprA* gene, known to be involved in oxidative stress resistance (Herman et al., 2016), was upregulated by 3.18-fold.

**Figure 6.**
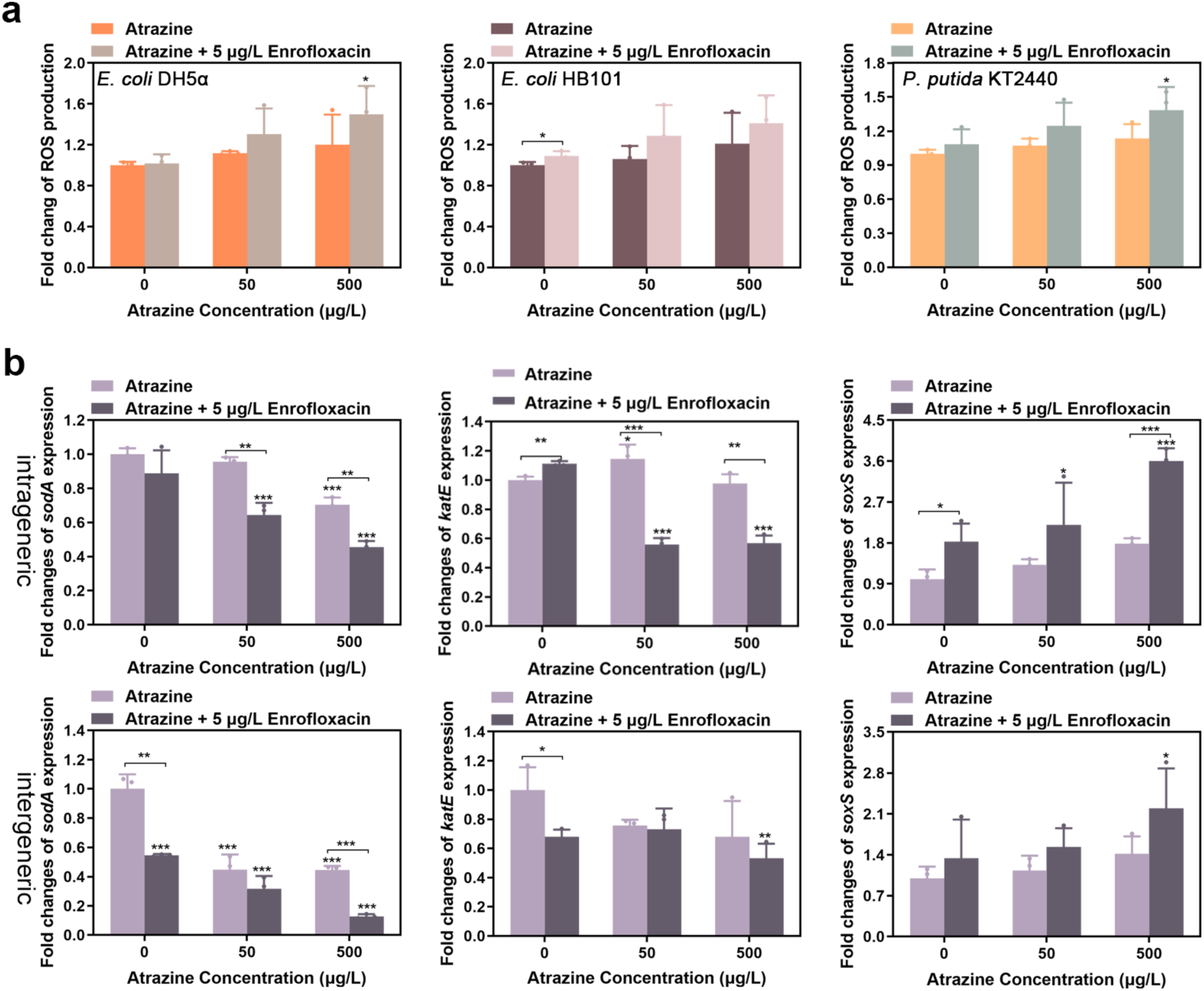
(a) The effects of atrazine alone or combined with enrofloxacin on ROS production in *E. coli* DH5α, *E. coli* HB101, *P. putida* KT2440. (b) Expression of *soxS*, *sodA* and *katE* genes under the exposure atrazine alone or in combination with enrofloxacin.

Furthermore, qPCR analysis revealed a 2.19-to 3.60-fold increase in the expression of the superoxide stress gene *soxS* in intra- and inter-genus conjugation systems, respectively, under combined exposure (Figure 6b) (Li and Demple, 1994; Xie et al., 2019). In contrast, genes encoding antioxidant enzymes, including *sodA* and *katE*, were significantly downregulated by 0.13–0.46-fold and 0.53–0.57-fold in intra- and inter-genus conjugation systems, respectively. These results indicate that the co-exposure disrupted the balance of the cellular redox system and exceeded the antioxidant capacity of the cells (Wang et al., 2020b). To further test the role of oxidative stress, ROS scavengers were introduced into the conjugation system. Their addition reduced the conjugation frequency of the co-exposure group, although the frequency remained higher than that of the control (Figure S4). This finding suggests that while oxidative stress contributes to the promotion of conjugative transfer, it is not the sole mechanism, and multiple pathways are likely involved.

## Resource availability

### Lead contact

Further information and reasonable requests for resources and reagents should be directed to and will be fulfilled by the lead contact, Pengfei Cui.

### Materials availability

Requests for materials should be made via the lead contact. All unique/stable reagents generated in this study are available from the lead contact without restriction.

### Data and code availability

- ***Data***: All data reported in this paper will be shared by the lead contact [Pengfei Cui] upon reasonable requests.
- ***Code***: This paper does not report original code.
- ***Additional Information***: Any additional information required to reanalyze the data reported in this article is available from the lead contact upon request.

## Acknowledgments

This work was supported by National Natural Science Foundation of China (No. 31902421).

## References

Avalos Vizcarra, I., Hosseini, V., Kollmannsberger, P., Meier, S., Weber, S.S., Arnoldini, M., Ackermann, M., Vogel, V., 2016. How type 1 fimbriae help Escherichia coli to evade extracellular antibiotics. Sci Rep 6, 18109. 10.1038/srep18109

Baharoglu, Z., Bikard, D., Mazel, D., 2010. Conjugative DNA Transfer Induces the Bacterial SOS Response and Promotes Antibiotic Resistance Development through Integron Activation. PLoS Genet 6, e1001165. 10.1371/journal.pgen.1001165

Bergogne-Bérézin, E., Towner, K.J., 1996. Acinetobacter spp. as nosocomial pathogens: microbiological, clinical, and epidemiological features. CLIN. MICROBIOL. REV. 9.

Cen, T., Zhang, X., Xie, S., Li, D., 2020. Preservatives accelerate the horizontal transfer of plasmid-mediated antimicrobial resistance genes via differential mechanisms. Environment International 138, 105544. 10.1016/j.envint.2020.105544

Chen, I., Christie, P.J., Dubnau, D., 2005. The Ins and Outs of DNA Transfer in Bacteria. Science 310, 1456–1460. 10.1126/science.1114021

Cheng, Q., Liu, Y., Xu, L., Ye, J., Wang, Q., Lin, H., Ma, J., 2023. Regulation and role of extracellular polymeric substances in the defensive responses of Dictyosphaerium sp. to enrofloxacin stress. Science of The Total Environment 896, 165302. 10.1016/j.scitotenv.2023.165302

Christie, P.J., Whitaker, N., González-Rivera, C., 2014. Mechanism and structure of the bacterial type IV secretion systems. Biochimica et Biophysica Acta (BBA) - Molecular Cell Research 1843, 1578–1591. 10.1016/j.bbamcr.2013.12.019

Crane, J.M., Randall, L.L., 2017. The Sec System: Protein Export in *Escherichia coli*. EcoSal Plus 7, ecosalplus.ESP-0002-2017. 10.1128/ecosalplus.ESP-0002-2017

Errington, J., Bath, J., Wu, L.J., 2001. DNA transport in bacteria. Nat Rev Mol Cell Biol 2, 538–545. 10.1038/35080005

Gao, B., Liang, L., Su, L., Wen, A., Zhou, C., Feng, Y., 2023. Structural basis for regulation of SOS response in bacteria. Proc. Natl. Acad. Sci. U.S.A. 120, e2217493120. 10.1073/pnas.2217493120

Garvey, N., St John, A.C., Witkin, E.M., 1985. Evidence for RecA protein association with the cell membrane and for changes in the levels of major outer membrane proteins in SOS-induced Escherichia coli cells. J Bacteriol 163, 870–876. 10.1128/jb.163.3.870-876.1985

Gutiérrez, R., Ram, Y., Berman, J., De Sousa, K.C.M., Nachum-Biala, Y., Britzi, M., Elad, D., Glaser, G., Covo, S., Harrus, S., 2021. Adaptive Resistance Mutations at Suprainhibitory Concentrations Independent of SOS Mutagenesis. Molecular Biology and Evolution 38, 4095–4115. 10.1093/molbev/msab196

Herman, A., Serfecz, J., Kinnally, A., Crosby, K., Youngman, M., Wykoff, D., Wilson, J.W., 2016. The Bacterial *iprA* Gene Is Conserved across Enterobacteriaceae, Is Involved in Oxidative Stress Resistance, and Influences Gene Expression in Salmonella enterica Serovar Typhimurium. J Bacteriol 198, 2166–2179. 10.1128/JB.00144-16

Holt, H.M., Gahrn-Hansen, B., Bruun, B., 2005. Shewanella algae and Shewanella putrefaciens: clinical and microbiological characteristics. Clinical Microbiology and Infection 11, 347–352. 10.1111/j.1469-0691.2005.01108.x

Horstmann, J.A., Lunelli, M., Cazzola, H., Heidemann, J., Kühne, C., Steffen, P., Szefs, S., Rossi, C., Lokareddy, R.K., Wang, C., Lemaire, L., Hughes, K.T., Uetrecht, C., Schlüter, H., Grassl, G.A., Stradal, T.E.B., Rossez, Y., Kolbe, M., Erhardt, M., 2020. Methylation of Salmonella Typhimurium flagella promotes bacterial adhesion and host cell invasion. Nat Commun 11, 2013. 10.1038/s41467-020-15738-3

Hu, Q., Zhang, L., Yang, R., Tang, J., Dong, G., 2024. Quaternary ammonium biocides promote conjugative transfer of antibiotic resistance gene in structure- and species-dependent manner. Environ. Int. 189, 108812. 10.1016/j.envint.2024.108812

Huang, H., Feng, G., Wang, M., Liu, C., Wu, Y., Dong, L., Feng, L., Zheng, X., Chen, Y., 2022. Nitric Oxide: A Neglected Driver for the Conjugative Transfer of Antibiotic Resistance Genes among Wastewater Microbiota. Environ. Sci. Technol. 56, 6466–6478. 10.1021/acs.est.2c01889

Ji, H., Cai, Y., Wang, Z., Li, G., An, T., 2022. Sub-lethal photocatalysis promotes horizontal transfer of antibiotic resistance genes by conjugation and transformability. Water Research 221, 118808. 10.1016/j.watres.2022.118808

Konig, B., Muller, J.J., Lanka, E., Heinemann, U., 2009. Crystal structure of KorA bound to operator DNA: insight into repressor cooperation in RP4 gene regulation. Nucleic Acids Research 37, 1915–1924. 10.1093/nar/gkp044

Li, X., Wen, C., Liu, C., Lu, S., Xu, Z., Yang, Q., Chen, Z., Liao, H., Zhou, S., 2022. Herbicide promotes the conjugative transfer of multi-resistance genes by facilitating cellular contact and plasmid transfer. Journal of Environmental Sciences 115, 363–373. 10.1016/j.jes.2021.08.006

Li, Z., Demple, B., 1994. SoxS, an activator of superoxide stress genes in Escherichia coli. Purification and interaction with DNA. Journal of Biological Chemistry 269, 18371–18377. 10.1016/S0021-9258(17)32317-7

Liao, H., Li, X., Yang, Q., Bai, Y., Cui, P., Wen, C., Liu, C., Chen, Z., Tang, J., Che, J., Yu, Z., Geisen, S., Zhou, S., Friman, V.-P., Zhu, Y.-G., 2021. Herbicide Selection Promotes Antibiotic Resistance in Soil Microbiomes. Molecular Biology and Evolution 38, 2337–2350. 10.1093/molbev/msab029

Liao, J., Huang, H., Chen, Y., 2019. CO2 promotes the conjugative transfer of multiresistance genes by facilitating cellular contact and plasmid transfer. Environment International 129, 333–342. 10.1016/j.envint.2019.05.060

Liu, Q., Wang, L., Chen, H., Huang, B., Xu, J., Li, Y., Héroux, P., Zhu, X., Wu, Y., Xia, D., 2019. Prometryn induces apoptotic cell death through cell cycle arrest and oxidative DNA damage. Toxicology Research 8, 833–841. 10.1039/c9tx00080a

Liu, X., Wang, D., Tang, J., Liu, F., Wang, L., 2021. Effect of dissolved biochar on the transfer of antibiotic resistance genes between bacteria. Environmental Pollution 288, 117718. 10.1016/j.envpol.2021.117718

Lu, J., Wang, Y., Li, J., Mao, L., Nguyen, S.H., Duarte, T., Coin, L., Bond, P., Yuan, Z., Guo, J., 2018. Triclosan at environmentally relevant concentrations promotes horizontal transfer of multidrug resistance genes within and across bacterial genera. Environment International 121, 1217–1226. 10.1016/j.envint.2018.10.040

Ma, X., Zhang, Xiuwen, Xia, J., Sun, H., Zhang, Xuxiang, Ye, L., 2021. Phenolic compounds promote the horizontal transfer of antibiotic resistance genes in activated sludge. Science of The Total Environment 800, 149549. 10.1016/j.scitotenv.2021.149549

Mittler, R., 2002. Oxidative stress, antioxidants and stress tolerance. Trends in Plant Science 7, 405–410. 10.1016/S1360-1385(02)02312-9

More, T.T., Yadav, J.S.S., Yan, S., Tyagi, R.D., Surampalli, R.Y., 2014. Extracellular polymeric substances of bacteria and their potential environmental applications. Journal of Environmental Management 144, 1–25. 10.1016/j.jenvman.2014.05.010

Ollis, A.A., Manning, M., Held, K.G., Postle, K., 2009. Cytoplasmic membrane protonmotive force energizes periplasmic interactions between ExbD and TonB. Molecular Microbiology 73, 466–481. 10.1111/j.1365-2958.2009.06785.x

O’neill, J. (2014) Antimicrobial Resistance: tackling a crisis for the health and wealth of nations.

Palmer, T., Berks, B.C., 2012. The twin-arginine translocation (Tat) protein export pathway. Nat Rev Microbiol 10, 483–496. 10.1038/nrmicro2814

Pu, Q., Fan, X.-T., Li, H., An, X.-L., Lassen, S.B., Su, J.-Q., 2021. Cadmium enhances conjugative plasmid transfer to a fresh water microbial community. Environmental Pollution 268, 115903. 10.1016/j.envpol.2020.115903

Qiu, Z., Yu, Y., Chen, Z., Jin, M., Yang, D., Zhao, Z., Wang, J., Shen, Z., Wang, X., Qian, D., Huang, A., Zhang, B., Li, J.-W., 2012. Nanoalumina promotes the horizontal transfer of multiresistance genes mediated by plasmids across genera. Proc. Natl. Acad. Sci. U.S.A. 109, 4944–4949. 10.1073/pnas.1107254109

Quiñones, A., Kaasch, J., Kaasch, M., Messer, W., 1989. Induction of dnaN and dnaQ gene expression in Escherichia coli by alkylation damage to DNA. The EMBO Journal 8, 587–593. 10.1002/j.1460-2075.1989.tb03413.x

Samuels, A.L., Lanka, E., Davies, J.E., 2000. Conjugative Junctions in RP4-Mediated Mating of *Escherichia coli*. J Bacteriol 182, 2709–2715. 10.1128/JB.182.10.2709-2715.2000

Silale, A., Van Den Berg, B., 2023. TonB-Dependent Transport Across the Bacterial Outer Membrane. Annu. Rev. Microbiol. 77, annurev-micro-032421-111116. 10.1146/annurev-micro-032421-111116

Stara, A., Kristan, J., Zuskova, E., Velisek, J., 2013. Effect of chronic exposure to prometryne on oxidative stress and antioxidant response in common carp (Cyprinus carpio L.). Pesticide Biochemistry and Physiology 105, 18–23. 10.1016/j.pestbp.2012.11.002

Su, C.-C., Yang, F., Long, F., Reyon, D., Routh, M.D., Kuo, D.W., Mokhtari, A.K., Van Ornam, J.D., Rabe, K.L., Hoy, J.A., Lee, Y.J., Rajashankar, K.R., Yu, E.W., 2009. Crystal Structure of the Membrane Fusion Protein CusB from Escherichia coli. Journal of Molecular Biology 393, 342–355. 10.1016/j.jmb.2009.08.029

Tan, Q., Li, W., Zhang, J., Zhou, W., Chen, J., Li, Y., Ma, J., 2019. Presence, dissemination and removal of antibiotic resistant bacteria and antibiotic resistance genes in urban drinking water system: A review. Front. Environ. Sci. Eng. 13, 36. 10.1007/s11783-019-1120-9

Thomas, C.M., Nielsen, K.M., 2005. Mechanisms of, and Barriers to, Horizontal Gene Transfer between Bacteria. Nat Rev Microbiol 3, 711–721. 10.1038/nrmicro1234

Tsirigotaki, A., De Geyter, J., Šoštarić, N., Economou, A., Karamanou, S., 2017. Protein export through the bacterial Sec pathway. Nat Rev Microbiol 15, 21–36. 10.1038/nrmicro.2016.161

Tsuneda, S., Aikawa, H., Hayashi, H., Yuasa, A., Hirata, A., 2003. Extracellular polymeric substances responsible for bacterial adhesion onto solid surface. FEMS Microbiology Letters 223, 287–292. 10.1016/S0378-1097(03)00399-9

UNEP 2017 Frontiers 2017 Emerging Issues of Environmental Concern. Solheim, E. (ed), Nairobi.

von Wintersdorff, C.J.H., Penders, J., van Niekerk, J.M., Mills, N.D., Majumder, S., van Alphen, L.B., Savelkoul, P.H.M., Wolffs, P.F.G., 2016. Dissemination of Antimicrobial Resistance in Microbial Ecosystems through Horizontal Gene Transfer. Front. Microbiol. 7. 10.3389/fmicb.2016.00173

Wang, J., Ding, Q., Yang, Q., Fan, H., Yu, G., Liu, F., Bello, B.K., Zhang, X., Zhang, T., Dong, J., Liu, G., Zhao, P., 2021. Vibrio alginolyticus Triggers Inflammatory Response in Mouse Peritoneal Macrophages via Activation of NLRP3 Inflammasome. Front. Cell. Infect. Microbiol. 11, 769777. 10.3389/fcimb.2021.769777

Wang, Q., Liu, L., Hou, Z., Wang, L., Ma, D., Yang, G., Guo, S., Luo, J., Qi, L., Luo, Y., 2020a. Heavy metal copper accelerates the conjugative transfer of antibiotic resistance genes in freshwater microcosms. Science of The Total Environment 717, 137055. 10.1016/j.scitotenv.2020.137055

Wang, X., Hu, M., Gu, H., Zhang, L., Shang, Y., Wang, Ting, Wang, Tingyue, Zeng, J., Ma, L., Huang, W., Wang, Y., 2020b. Short-term exposure to norfloxacin induces oxidative stress, neurotoxicity and microbiota alteration in juvenile large yellow croaker Pseudosciaena crocea. Environmental Pollution 267, 115397. 10.1016/j.envpol.2020.115397

Wang, Y., Lu, J., Mao, L., Li, J., Yuan, Z., Bond, P.L., Guo, J., 2019. Antiepileptic drug carbamazepine promotes horizontal transfer of plasmid-borne multi-antibiotic resistance genes within and across bacterial genera. ISME J 13, 509–522. 10.1038/s41396-018-0275-x

Wang, Z., Sun, X., Ru, S., Wang, J., Xiong, J., Yang, L., Hao, L., Zhang, J., Zhang, X., 2022. Effects of co-exposure of the triazine herbicides atrazine, prometryn and terbutryn on Phaeodactylum tricornutum photosynthesis and nutritional value. Science of The Total Environment 807, 150609. 10.1016/j.scitotenv.2021.150609

Xie, S., Gu, A.Z., Cen, T., Li, D., Chen, J., 2019. The effect and mechanism of urban fine particulate matter (PM2.5) on horizontal transfer of plasmid-mediated antimicrobial resistance genes. Science of The Total Environment 683, 116–123. 10.1016/j.scitotenv.2019.05.115

Yang, B., Wang, Z., Jia, Y., Fang, D., Li, R., Liu, Y., 2022. Paclitaxel and its derivative facilitate the transmission of plasmid-mediated antibiotic resistance genes through conjugative transfer. Science of The Total Environment 810, 152245. 10.1016/j.scitotenv.2021.152245

Yang, L., He, X., Ru, S., Zhang, Y., 2024. Herbicide leakage into seawater impacts primary productivity and zooplankton globally. Nat Commun 15, 1783. 10.1038/s41467-024-46059-4

Yang, L., Li, H., Zhang, Y., Jiao, N., 2019. Environmental risk assessment of triazine herbicides in the Bohai Sea and the Yellow Sea and their toxicity to phytoplankton at environmental concentrations. Environment International 133, 105175. 10.1016/j.envint.2019.105175

Yu, K., Chen, F., Yue, L., Luo, Y., Wang, Z., Xing, B., 2020. CeO_2_ Nanoparticles Regulate the Propagation of Antibiotic Resistance Genes by Altering Cellular Contact and Plasmid Transfer. Environ. Sci. Technol. 54, 10012–10021. 10.1021/acs.est.0c01870

Yu, Z., Wang, Y., Lu, J., Bond, P.L., Guo, J., 2021. Nonnutritive sweeteners can promote the dissemination of antibiotic resistance through conjugative gene transfer. ISME J 15, 2117–2130. 10.1038/s41396-021-00909-x

Zatyka, M., Bingle, L., Jones, A.C., Thomas, C.M., 2001. Cooperativity between KorB and TrbA Repressors of Broad-Host-Range Plasmid RK2. J Bacteriol 183, 1022–1031. 10.1128/JB.183.3.1022-1031.2001

Zatyka, M., Jagura-Burdzy, G., Thomas, C.M., 1997. Transcriptional and translational control of the genes for the mating pair formation apparatus of promiscuous IncP plasmids. J Bacteriol 179, 7201–7209. 10.1128/jb.179.23.7201-7209.1997

Zhang, H., Song, J., Zheng, Z., Li, T., Shi, N., Han, Y., Zhang, L., Yu, Y., Fang, H., 2023. Fungicide exposure accelerated horizontal transfer of antibiotic resistance genes via plasmid-mediated conjugation. Water Res. 233, 119789. 10.1016/j.watres.2023.119789

Zhang, H., Xu, L., Hou, X., Li, Y., Niu, L., Zhang, J., Wang, X., 2024a. Ketoprofen promotes the conjugative transfer of antibiotic resistance among antibiotic resistant bacteria in natural aqueous environments. Environ. Pollut. 360, 124676. 10.1016/j.envpol.2024.124676

Zhang, S., Wang, Y., Song, H., Lu, J., Yuan, Z., Guo, J., 2019. Copper nanoparticles and copper ions promote horizontal transfer of plasmid-mediated multi-antibiotic resistance genes across bacterial genera. Environment International 129, 478–487. 10.1016/j.envint.2019.05.054

Zhang, Y., Gu, A.Z., Cen, T., Li, X., He, M., Li, D., Chen, J., 2018. Sub-inhibitory concentrations of heavy metals facilitate the horizontal transfer of plasmid-mediated antibiotic resistance genes in water environment. Environmental Pollution 237, 74–82. 10.1016/j.envpol.2018.01.032

Zhang, Y., Meng, D., Wang, Z., Guo, H., Wang, Y., 2012. Oxidative stress response in two representative bacteria exposed to atrazine. FEMS Microbiol Lett 334, 95–101. 10.1111/j.1574-6968.2012.02625.x

Zhang, Z., Feng, Y., Wang, W., Ru, S., Zhao, L., Ma, Y., Song, X., Liu, L., Wang, J., 2024b. Pollution level and ecological risk assessment of triazine herbicides in laizhou bay and derivation of seawater quality criteria. J. Hazard. Mater. 477, 135270. 10.1016/j.jhazmat.2024.135270

Zhang, Z., Zhang, Q., Wang, T., Xu, N., Lu, T., Hong, W., Penuelas, J., Gillings, M., Wang, M., Gao, W., Qian, H., 2022. Assessment of global health risk of antibiotic resistance genes. Nat. Commun. 13, 1553. 10.1038/s41467-022-29283-8

Zhao, Q., Huang, M., Yin, J., Wan, Y., Liu, Y., Duan, R., Luo, Y., Xu, X., Cao, X., Yi, M., 2022b. Atrazine exposure and recovery alter the intestinal structure, bacterial composition and intestinal metabolites of male Pelophylax nigromaculatus. Science of The Total Environment 818, 151701. 10.1016/j.scitotenv.2021.151701

Zhao, R., Jiang, S., Zhang, L., Yu, Z., 2019. Mitochondrial electron transport chain, ROS generation and uncoupling (Review). Int J Mol Med. 10.3892/ijmm.2019.4188

Zhao, Y., Cao, Z., Cui, L., Hu, T., Guo, K., Zhang, F., Wang, X., Peng, Z., Liu, Q., Dai, M., 2022a. Enrofloxacin promotes plasmid-mediated conjugation transfer of fluoroquinolone-resistance gene *qnrS*. Front. Microbiol. 12, 773664. 10.3389/fmicb.2021.773664

